# Speech intelligibility predicted from neural entrainment of the speech envelope

**DOI:** 10.1101/246660

**Authors:** Jonas Vanthornhout, Lien Decruy, Jan Wouters, Jonathan Z. Simon, Tom Francart

## Abstract

Speech intelligibility is currently measured by scoring how well a person can identify a speech signal. The results of such behavioral measures reflect neural processing of the speech signal, but are also influenced by language processing, motivation and memory. Very often electrophysiological measures of hearing give insight in the neural processing of sound. However, in most methods non-speech stimuli are used, making it hard to relate the results to behavioral measures of speech intelligibility. The use of natural running speech as a stimulus in electrophysiological measures of hearing is a paradigm shift which allows to bridge the gap between behavioral and electrophysiological measures. Here, by decoding the speech envelope from the electroencephalogram, and correlating it with the stimulus envelope, we demonstrate an electrophysiological measure of neural processing of running speech. We show that behaviorally measured speech intelligibility is strongly correlated with our electrophysiological measure. Our results pave the way towards an objective and automatic way of assessing neural processing of speech presented through auditory prostheses, reducing confounds such as attention and cognitive capabilities. We anticipate that our electrophysiological measure will allow better differential diagnosis of the auditory system, and will allow the development of closed-loop auditory prostheses that automatically adapt to individual users.

## 1 Introduction

The human auditory system processes speech in different stages. The auditory periphery converts the sound pressure wave into neural spike trains, the auditory cortex segregates streams, and finally specialized language processing areas are activated, which interact with short and long term memory. Each of these subsystems can be impaired, so in diagnostics it is crucial to be able to measure the function of the auditory system at the different levels. The current audiometric test battery consists of behavioral tests of speech intelligibility and objective measures based on electroencephalogram (EEG).

In behavioral tests of speech intelligibility the function of the entire auditory system is measured. A fragment of natural speech is presented and the subject is instructed to identify it. When the goal is to assess the function of the auditory periphery, such as fitting auditory prostheses, language knowledge and cognitive function such as working memory are confounds. Additionally, behavioral testing requires active participation of the test subject, which is not always possible and leads to another confound: motivation and attention. With current EEG-based objective measures, it is possible to measure the function of intermediate stages of the auditory system, but unnatural periodic stimuli, such as click trains, modulated tones or repeated phonemes are used (e.g., Anderson et al, 2013; Picton et al, 2005; McGee and Clemis, 1980), which are acoustically different from natural running speech, and are processed differently by the brain (Hullett et al, 2016). While these measures yield valuable information about the auditory system, they are not well-correlated with behaviorally measured speech intelligibility. Another practical downside of non-speech stimuli is that they may be processed differently from speech by modern auditory prostheses which take into account the statistics of speech signals (Dillon, 2012). This is problematic when assessing a subject’s hearing through an auditory prosthesis such as a hearing aid or cochlear implant.

The missing link between behavioral and objective measures is a measure of neural processing of the acoustic cues in speech that lead to intelligibility. The most important acoustic cue for speech intelligibility is the temporal envelope (Shannon et al, 1995; Peelle and Davis, 2012) and especially modulation frequencies below 20 Hz (Drullman et al, 1994b,a). Recently, it has been shown with non-invasive magnetoencephalography (MEG) and EEG recordings that neural processing of the speech envelope can be inferred from the correlation between the actual speech envelope and the speech envelope decoded from the neural signal (Aiken and Picton, 2008; Ding and Simon, 2011). Even for running speech in a single-trial paradigm i.e., presenting the stimulus only once the speech envelope could reliably be reconstructed (Ding and Simon, 2012, 2013; O’Sullivan et al, 2014; Di Liberto et al, 2015; Horton et al, 2014). A decoder transforms the multi-channel neural signal into a single-channel speech envelope, by linearly combining amplitude samples across MEG sensors and across a post-stimulus temporal integration window. Based on training data, the decoder is calculated as the linear combination that maximizes the correlation with the actual speech envelope. This method has also been shown to work with electroencephalography (EEG) recordings (O’Sullivan et al, 2014). Furthermore, using surface recordings of the cortex, the full stimulus spectrogram can be decoded (Pasley et al, 2012), and inversely the full spectrogram and even phoneme representation can be used to predict the EEG signal (Di Liberto et al, 2015).

Using these techniques, previous research has compared the correlation between the speech envelope and the reconstructed envelope, with speech intelligibility (Ding and Simon, 2013; Kong et al, 2015). However, the interpretation of the results is complicated by the fact that speech intelligibility could fluctuate over time due to the use of non-standardized running speech as a stimulus, and because subjective ratings were used as a measure of speech intelligibility instead of standardized speech audiometry. Standardized audiometric speech materials are carefully optimized for precision and reliability, something which is difficult, if not impossible with running speech and subjective ratings.

Therefore, we developed an objective measure of neural processing of the speech envelope based on the stimulus reconstruction method and compared it with behaviorally measured speech intelligibility. We do not expect these measures to correspond exactly, as there are some inherent differences, in particular the higher level functions such as working memory and cognitive function that are relied upon for the behavioural measure and not so much for the objective one. However, on the one hand we reduced those differences by the choice of materials and methods, and on othe other hand it remains important to compare our novel objective measure to the current gold standard for measuring speech intelligibility. We used EEG rather than MEG, as it is ubiquitous, can be implemented on a large scale, and is often available for clinical application.

## 2 Methods

An overview of our methods is shown in Figure 1. Briefly, in a behavioral and EEG experiment, we used the same speech stimuli, from a standardized speech test, combined with spectrally matched stationary noise at different signal to noise ratios (SNRs). In the behavioral experiment, we determined the speech reception threshold (SRT). In the EEG experiment, we determined neural entrainment of the speech envelope as a function of SNR, and derived an objective measure. We then compared the SRT with the objective measure on an individual subject basis.

**Figure 1:**
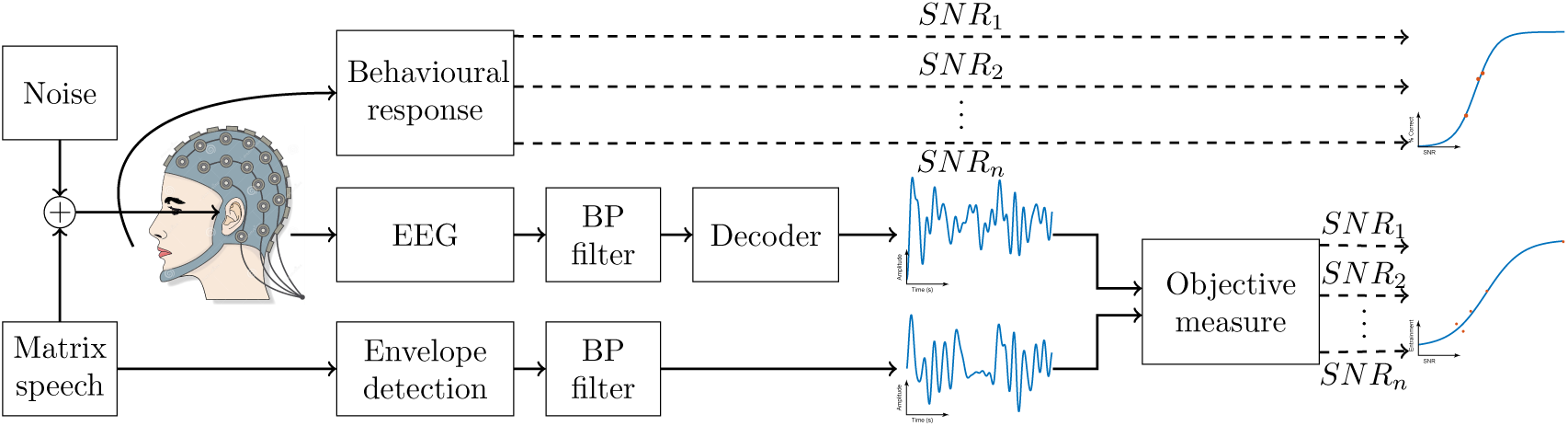
Overview of the experimental setup. We used the Flemish Matrix sentences to behaviorally measure speech intelligibility. In the EEG experiment we presented stimuli from the same Matrix corpus while measuring the EEG. By correlating the speech envelopes from the Matrix and the envelopes decoded from the EEG, we obtained our objective measure.

The objective measure is obtained by on the one hand determining the slowly varying temporal envelope of the speech signal (bottom row of Figure 1), which can be thought of as the signal power over time, and on the other hand attempting to decode this same envelope from the EEG signal (middle row of Figure 1). To this end, for each subject a decoder is trained on speech in quiet, which decodes the speech envelope as a linear combination of EEG samples, across a temporal integration window, and across the EEG recording electrodes. The actual and decoded envelopes are then correlated with each other, which yields a measure of neural entrainment of the speech envelope. After repeating this process for a number of SNRs, a sigmoid function is fitted to the results. The midpoint of the resulting sigmoid function is our objective measure, which we call the correlation threshold (CT).

**Figure 2:**
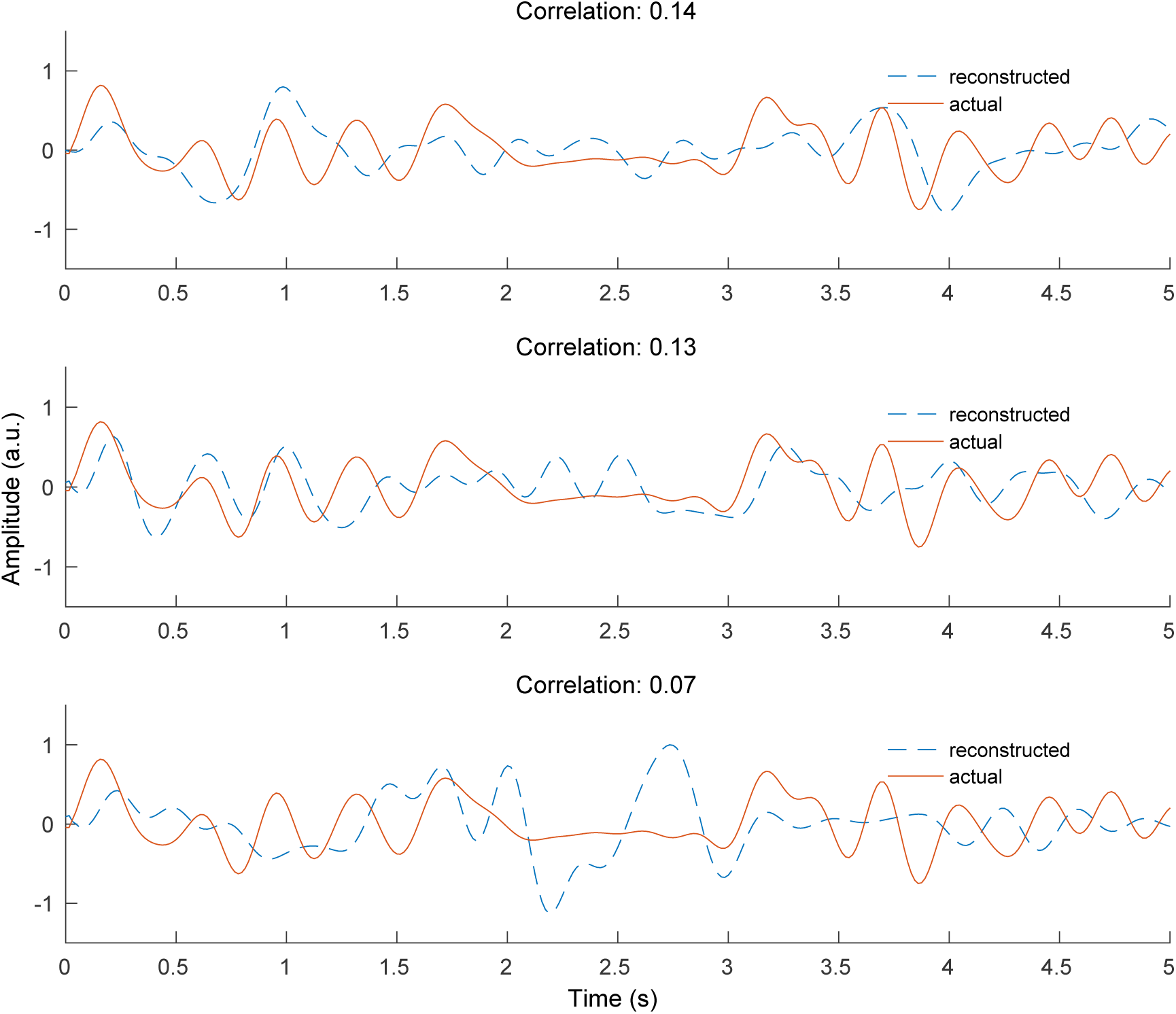
Examples of actual and reconstructed envelopes and the corresponding correlations.

### 2.1 Participants

We tested 24 normal-hearing subjects, 7 male and 17 female, recruited from our university student population to ensure normal language processing and cognitive function. Their age ranged from 21 to 29 years with an average of 24.3 years. Every subject reported normal hearing, which was verified by pure tone audiometry (thresholds lower than 25 dB HL for 125 Hz until 8000 Hz using MADSEN Orbiter 922-2). They had Dutch (Flemish) as mother tongue and were unpaid volunteers. Before each experiment the subjects signed an informed consent form approved by the Medical Ethics Committee UZ KU Leuven / Research (KU Leuven) with reference S59040.

### 2.2 Behavioral experiments

The behavioral experiments consisted of tests with the Flemish Matrix material Luts et al (2015) using the method of constant stimuli at 3 SNRs around the SRT. This material is divided in lists of 20 sentences which have been shown to yield similar behavioral speech intelligibility scores. Such validated tests, consisting of a standardized corpus of sentences, are currently the gold standard in measuring speech intelligibility, both in research and clinical practice. Sentences were spoken by a female speaker and presented to the right ear. They have a fixed structure of ‘name verb numeral adjective object’, where each element is selected from a closed set of ten possibilities, e.g., ‘Sofie ziet zes grijze pennen’ (‘Sofie sees six gray pens’). These sentences sound perfectly natural, but are grammatically trivial and completely unpredictable, thus minimizing the effect of higher order language processing.

The experiments were conducted on a laptop running Windows using the APEX 3 (version 3.1) software platform developed at ExpORL (Dept. Neurosciences, KU Leuven) (Francart et al, 2008), an RME Multiface II sound card (RME, Haimhausen, Germany) and Etymotic ER-3A insert phones (Etymotic Research, Inc., Illinois, USA) which were electromagnetically shielded using CFL2 boxes from Perancea Ltd. (London, United Kingdom). The speech was presented monaurally at 60 dBA and the setup was calibrated in a 2-cm^3^ coupler (Brüel & Kjaer 4152) using the stationary speech weighted noise corresponding with the Matrix speech material. The experiments took place in an electromagnetically shielded and soundproofed room.

### 2.3 EEG experiments

#### Setup

To measure auditory evoked potentials we used a BioSemi (Amsterdam, Netherlands) ActiveTwo EEG setup with 64 electrodes and recorded the data at a sampling rate of 8192 Hz using the ActiView software provided by BioSemi. The stimuli were presented with the same setup as the behavioral experiments, with the exception of diotic stimulation and adapting the noise level instead of the speech level for the EEG experiment.

#### Speech material

We presented stimuli created by concatenating two lists of Flemish Matrix sentences with a gap between the sentences. This length of this gap was uniformly distributed between 0.8 s and 1.2 s. The total duration of this stimulus was around 120 seconds. It was presented at 3, 5 or 7 different SNRs with the speech level fixed at 60 dBA. The order of SNRs was randomised across subjects. Each stimulus was presented 3 or 4 times. The total duration of the experiment was 2 hours. To keep the subjects attentive, questions about the stimuli were asked before and after the presentation of the stimulus. The questions were typically counting tasks, e.g. ‘How many times did you hear “gray pens”?’. These Matrix sentences were used to objectively estimate the speech understanding.

#### Speech story

The subjects listened to the children’s story ‘Milan’, written and narrated in Flemish by Stijn Vranken^1^. It was 15 minutes long and was presented at 60 dBA without any noise. The purpose of this stimulus was to have a continuous, attended stimulus to train the linear decoder. No questions were asked before or after this stimulus.

## 2.4 Signal processing

### Speech

We measured envelope entrainment by calculating the bootstrapped Spearman correlation (see below) between the stimulus speech envelope and the envelope reconstructed by a linear decoder. All implementations were written in MATLAB R2016b.

The stimulus speech envelope was extracted according to Biesmans et al (2016), who investigated the effect of envelope extraction method on auditory attention detection, and found best performance for a gammatone filterbank followed by a power law. In more detail, we used a gammatone filterbank (Søndergaard and Majdak, 2013; Søndergaard et al, 2012) with 28 channels spaced by 1 equivalent rectangular bandwidth (ERB), with center frequencies from 50 Hz until 5000 Hz. From each subband we extracted the envelope by taking the absolute value of each sample and raising it to the power of 0.6. The resulting 28 subband envelopes were averaged to obtain one single envelope. The power law was chosen as the human auditory system is not a linear system and compression is present in the system. The gammatone filterbank was chosen as it mimics the auditory filters present in the basilar membrane in the cochlea.

The speech envelope and EEG signal were band-pass filtered. We investigated performance for a range of filter cut-off frequencies. The same filter (a zero phase Butterworth filter with 80 dB attenuation at 10% outside the passband) was applied to the EEG and speech envelope. Before filtering, the EEG data were re-referenced to Cz and were downsampled from 8192 Hz to 1024 Hz to decrease processing time. After filtering, the data were further downsampled to 64 Hz.

A decoder, is a spatial filter, over EEG electrodes and a temporal filter, over time lags which optimally reconstructs the speech envelope from the EEG. The decoder linearly combines EEG electrode signals and their time shifted versions to optimally reconstruct the speech envelope. In the training phase, the weights to be applied to each signal in this linear combination are determined. The decoder was calculated using the mTRF toolbox (version 1.1) (Lalor et al, 2006, 2009) and applied as follows. As the stimulus evoked neural responses at different delays along the auditory pathway, we define a matrix *R* containing the shifted neural responses of each channel. If *g* is the linear decoder and *R* is the shifted neural data, the reconstruction of the speech envelope ŝ(*t*) was obtained as follows:

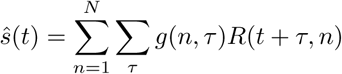

with *t* the time ranging from 0 to *T*, *n* the index of the recording electrode and *τ* the post-stimulus integration-window length used to reconstruct the envelope. The matrix *g* can be determined by minimizing a least-squares objective function

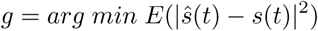

where *E* denotes the expected value, *s*(*t*) the real speech envelope and ŝ(*t*) the reconstructed envelope. In practice we calculated the decoder by solving

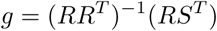

where *R* is the time-lagged matrix of the neural data and *S* a vector of stimulus envelope samples. The decoder is calculated using ridge regression on the inverse autocorrelation matrix.

We trained a new decoder for each subject on the story stimulus, which was 15 minutes long. After training, the decoder was applied on the EEG responses to the Flemish Matrix material.

To measure the correspondence between the speech envelope and its reconstruction, we calculated the bootstrapped Spearman correlation between the real and reconstructed envelope. Bootstrapping was applied by Monte Carlo sampling of the two envelopes. Some examples of actual and reconstructed envelopes and the corresponding correlations are shown in figure 2.

Our goal is to derive an objective measure of speech intelligibility, similar to the SRT for behavioral tests. Therefore the correlation between real and reconstructed envelope needs to increase with SNR, just like the percentage correctly repeated words increases with SNR in behavioral measures. To allow quantitative comparison between the different conditions of band pass filter and decoder temporal integration window, we defined a measure of monotonicity of the stimulus SNR versus correlation function. For each subject it indicates the percentage that the following comparisons are true: the correlation at the lowest SNR is lower than the correlations at the middle and highest SNR, and the correlation at the highest SNR is higher than the correlation at the lowest SNR. The band pass filter and temporal integration window were chosen to maximize this measure across all subjects.

## 3 Results

As different roles are attributed to different EEG frequency bands, we first investigated the effect of the cut-off frequencies of the band-pass filter that is applied to both the envelope and EEG signal. Next, we investigated the effect of integration window of the decoder. This can be understood as the number of EEG samples following the acoustic stimulus that are taken into account. For both the filter and the integration window we selected the parameter values that yielded optimal monotonicity of the entrainment versus SNR. Finally, using the optimal parameters, we calculated the correlation between the actual speech envelope and the reconstructed envelope for each SNR, derived our objective measure of speech intelligibility, and compared it to the behavioral SRT.

### 3.1 Filter band

Neural responses are mostly analyzed in specific filter bands. Much of the speech-related EEG research focuses on the delta band (0.5 Hz - 4 Hz) and theta band (4 Hz - 8 Hz) (O’Sullivan et al, 2014; Ding and Simon, 2013; Doelling et al, 2014). We systematically investigated the effect of low‐ and high-pass frequency of the band on monotonicity of the reconstruction quality as a function of stimulus SNR. We found best monotonicity using only the delta band (Figure 3a). Best performance was found when low frequencies are included. As a result we used a filter band from 0.5 until 4 Hz.

**Figure 3:**
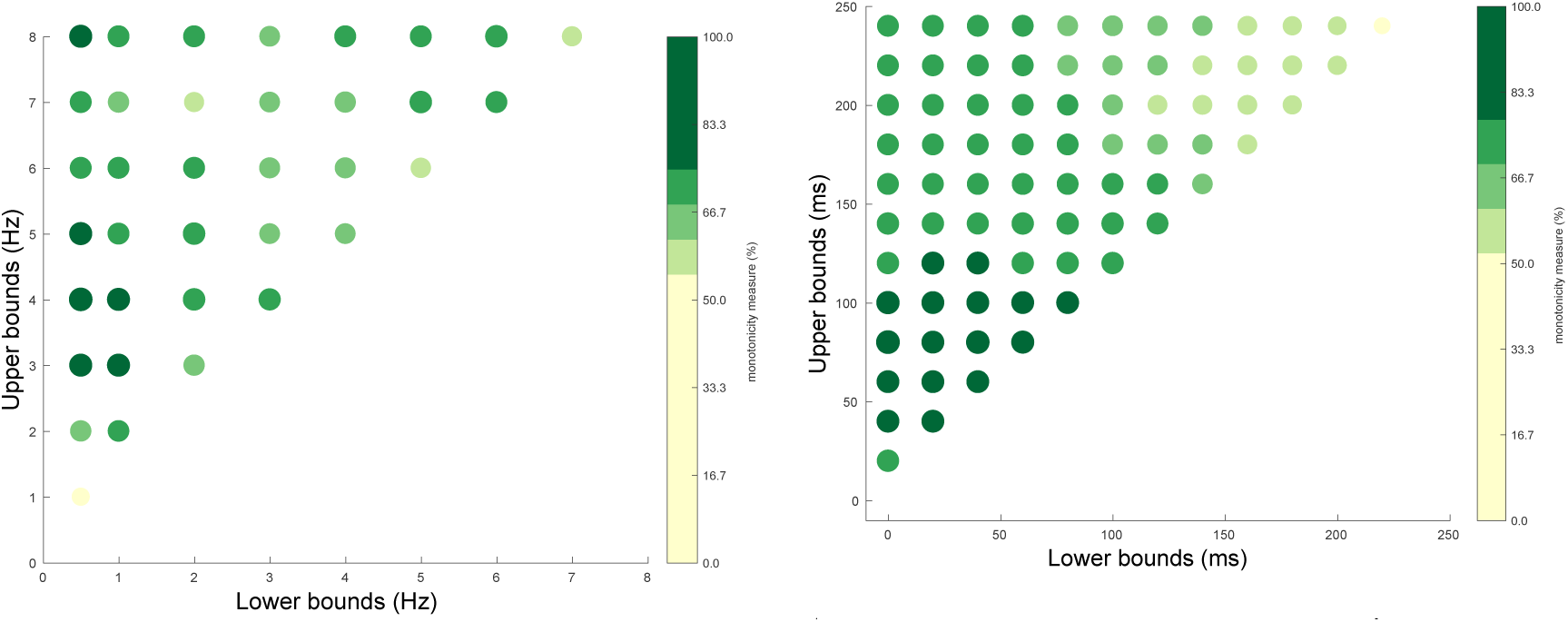
The monotonicity of envelope entrainment as a function of frequency bands and temporal integration window. (a) Monotonicity of envelope entrainment as a function of lower and upper bound of the pass band filter. Warm colors reflect a higher percentage correct. Best performance is seen when low frequencies (0.5 until 4 Hz) are included. (b) Monotonicity of envelope entrainment as a function of lower and upper bound of the temporal integration window of the decoder. Warm colors reflect a higher percentage correct. Best performance is seen for integration windows including early responses from 0 ms up to 75-140 ms.

### 3.2 Integration window

We systematically varied the temporal integration window of the decoder, and found best monotonicity of the reconstruction quality using an integration window focusing on early responses, from 0 ms up to 75-140 ms, see Figure 3b. Other research has shown that early responses yield a more gradual decline in correlation with decrease in SNR (Ding and Simon, 2013), compared to later responses, and that earlier responses are less modulated by attention (Ding and Simon, 2012; O’Sullivan et al, 2014). Based on these findings and our results, we used an integration window from 0 ms until 75 ms.

### 3.3 Behavioral versus Objective

Behavioral speech intelligibility was characterized by the speech reception threshold (SRT), i.e., the SNR yielding 50% intelligibility. It was obtained by fitting a sigmoid function with the formula 
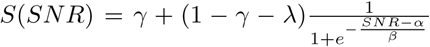
 with *γ* the guess-rate, λ the lapse-rate, *α* the midpoint and *β* the slope, to the SNR-versus-intelligibility points for each subject individually (e.g., Figure 4a). For the behavioral data, *γ* and were λ fixed to 0, leaving 2 parameters to be fitted to 3 data points, as is common for obtaining the SRT. The mean of the individual SRTs was −7.4 dB with an inter-subject standard deviation of 1.3 dB, ranging from −9.9 dB to −4.7 dB.

**Figure 4:**
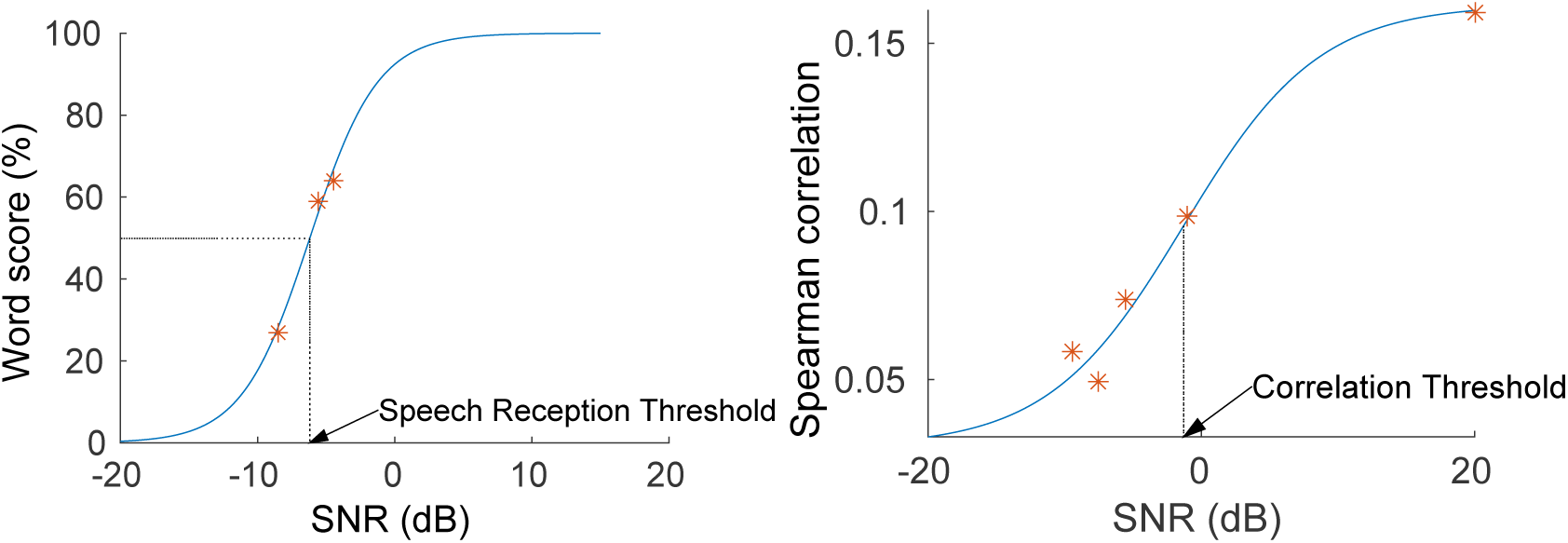
Behavioral and objective results for one subject. (a) The percentage of words correctly understood increases with increasing SNR. The blue line is a sigmoid function fitted on these data, from which we can estimate the speech reception threshold (SRT). (b) The Spearman correlation between actual speech envelope and speech envelope extracted from the EEG response increases with increasing SNR. The blue line is a sigmoid function fitted on these data, from which we can estimate our objective measure, the correlation threshold (CT).

The objective measure was inspired by the behavioral one in the sense that we obtained a single-trial score for each of a range of SNRs and then fitted a sigmoid function. The score was calculated as the absolute value of the Spearman correlation between the actual and the decoded speech envelope. In Figure 5 the scores for each subject and SNR are shown.

**Figure 5:**
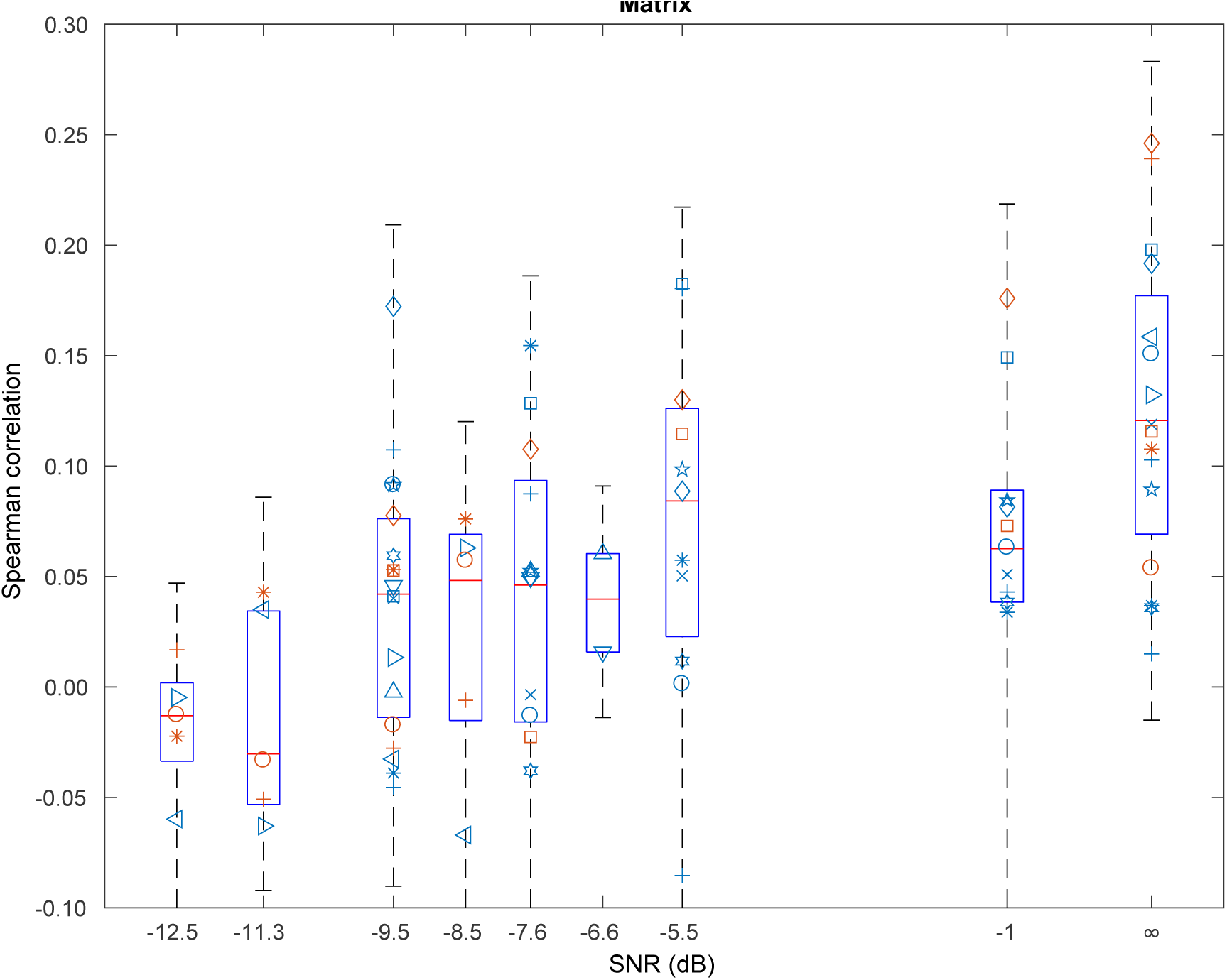
Individual data points of the entrainment over SNRs. Each subject is a different symbol, the boxplot gives an idea of the variance across subjects.

For the objective data, *γ* was fixed to 0:03, the chance level of the correlation. The chance level was computed by correlating the reconstructed envelope with a different part of the actual envelope. As a result we fitted the remaining 3 parameters to at least 5 data points. After fitting the function, we derived its midpoint, and used this as our objective measure, which we will refer to as the correlation threshold (CT), e.g., Figure 4b. The benefit of this measure, compared to using the correlation value at a single SNR directly, is that the target SNR, which is subject specific, does not need to be known a priori and that it is robust to inter-subject differences in correlation magnitude.

Using individual decoders we were able to obtain a good fit of the sigmoid function for 19 of the 24 subjects, i.e., no fitted parameter was equal to its lower or upper bound, and consequently derived the CT. We found a significant Pearson correlation of 0.69 between SRT and CT (p=0.001, Figure 6). Given the relatively small range of behavioral results for these normal-hearing subjects, from −9.9 dB SNR to −4.7 dB SNR, and a typical test-retest difference of 1 dB of the behavioral measure, this indicates that our objective measure is sensitive to small changes in SRT.

**Figure 6:**
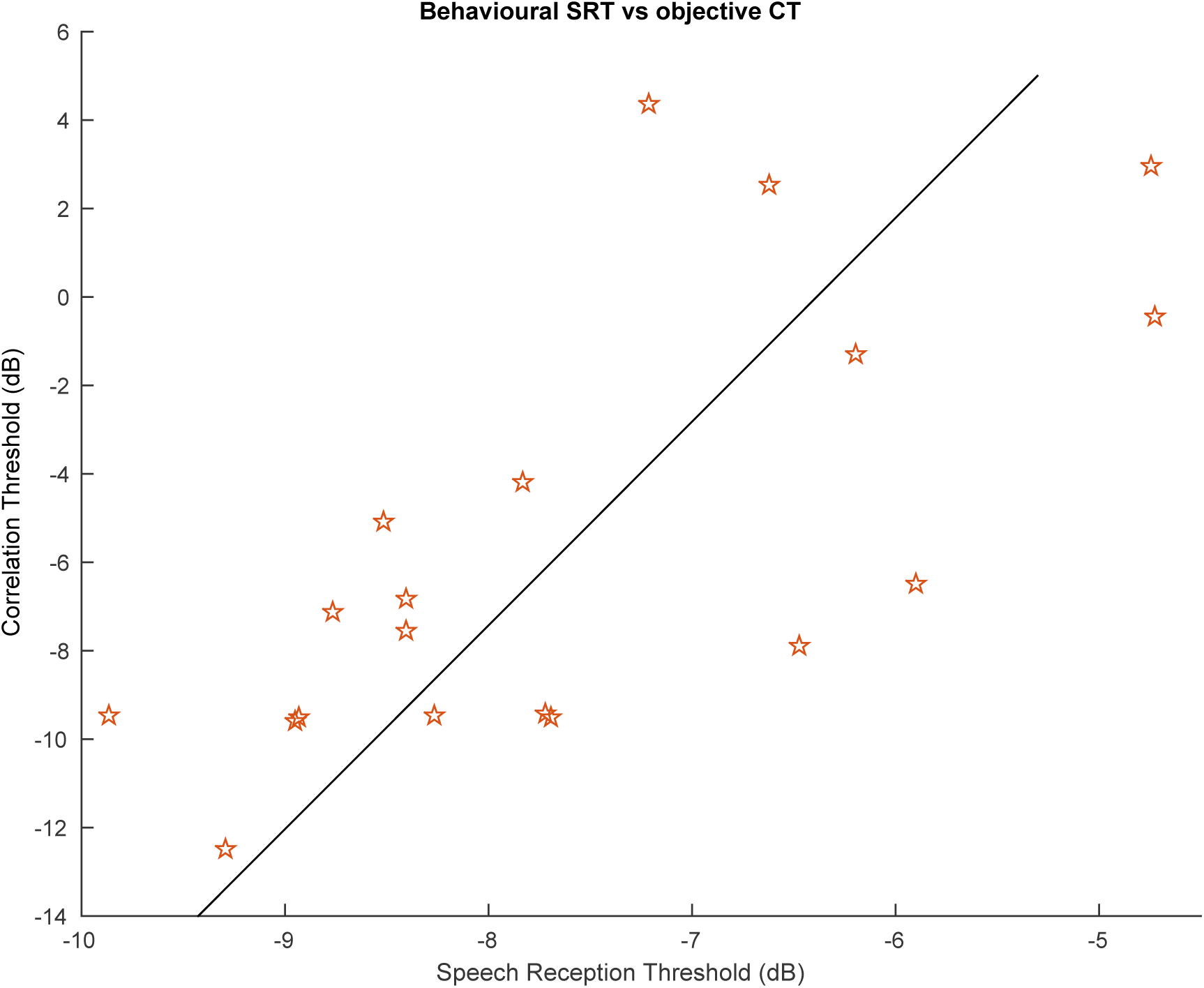
Electrophysiological versus behavioral measure (Pearson’s *r* = 0.69, *p* = 0.001). The electrophysiological measure (correlation threshold, CT) is the midpoint of each psychometric function. The behavioral measure (speech reception threshold, SRT) is the stimulus SNR at which the subject can understand 50% of the words.

## 4 Discussion

We compared a new objective measure of speech intelligibility (the CT) to the behaviorally measured SRT for 24 normal-hearing subjects. The objective measure is based on the correlation between the actual speech envelope and the speech envelope reconstructed from the EEG signal, a measure of neural entrainment to the speech envelope. We fitted a sigmoid function to the resulting entrainment versus stimulus SNR data, and derived the CT as its midpoint. We found a significant correlation between the objectively measured CT and behaviorally measured SRT.

### 4.1 Filter band

We found highest monotonicity in the delta band. This band encompasses the main information in the modulation spectrum of speech which exhibits peaks at the sentence rate (0.5 Hz) and word rate (2.5 Hz) (Edwards and Chang, 2013). It contains the prosodic information which is known to be important for speech intelligibility (Woodfield and Akeroyd, 2010). For the Matrix sentences, sharp peaks can be observed in the modulation spectrum at 0.5, 2.5 and 4.1 Hz, due to low variation among the sentences. Note that the delta band does not include the syllable rate of the Matrix sentences (4.1 Hz). Ding and Simon (2013); Ding et al (2014); Doelling et al (2014) also found that the neural responses in delta band were a predictor of how well individual subjects recognized speech in noise.

### 4.2 Integration window

We found best monotonicity of correlation as a function of SNR for an integration window from 0 ms until 75 ms. This may be counter-intuitive as higher correlation values, but not monotonicity are obtained using a longer integration window, such as 0 ms until 500 ms (Ding and Simon, 2013) and other studies focus more on later responses (O’Sullivan et al, 2014; Di Liberto et al, 2015). However recent work (Ding and Simon, 2012; O’Sullivan et al, 2014) shows that early responses (0 ms to 75 ms) are less modulated by attention compared to later responses (later than 75 ms). Our stimulus is unpredictable and not particularly engaging, so it is likely that the subjects were not attentive throughout the entire experiment (in spite of the instructions). By using only the early responses we limit the attentional effects.

### 4.3 Behavioral versus Objective

We found a significant correlation between the behaviorally measured SRT and our new objective measure (CT). Ding and Simon (2014) reviewed a number of studies in which similar comparisons are made. They concluded that in many cases stimuli which differ in intelligibility also differ in acoustic properties, making it difficult to determine if changes in cortical entrainment arise from changes in speech intelligibility or from changes in acoustic properties. We addressed this by using stimuli with similar statistics in all conditions. Additionally, in previous work, subjective ratings of intelligibility of a non-standardized story were used as the behavioral measurement. The problem is that such measures are prone to large inter-subject differences and larger variability than for standardized speech audiometry. We addressed this by using standardized speech material as the stimulus for both the behavioral and EEG experiments. Moreover, the correlation between actual and reconstructed envelope can differ widely in magnitude across subjects, due to differences in recording SNR of the EEG signal. Therefore we avoided using it directly and instead captured the trend across SNRs by fitting a sigmoid function.

Ding and Simon (2013) found a correlation between subjectively rated intelligibility and reconstruction accuracy in an MEG experiment. When assessing reconstruction accuracy as a function of SNR across subjects, they found that it was relatively unaffected down to a certain SNR and then sharply dropped. Possible explanations for the difference with our results, where we found a more gradual decrease in reconstruction accuracy with SNR, are the type of speech material used (low-context Matrix sentences versus a story) and the decoder integration window length (75 ms versus 250 ms).

The correlation between the SRT and the CT only explains 50 percent of the variance. The remainder can be attributed to limitations of our model, state of the subject, and limitations of the behavioural measure. In our model, we only used the speech envelope, which is a crude representation of a speech signal, and indeed the auditory system uses many other cues such as frequency-dependent envelopes and temporal fine structure. For instance, Di Liberto et al (2015) have shown that including the entire spectrogram or even a phoneme-representation of the stimulus can improve performance. Also, our simple linear decoder is probably not able to cope with all the complexity of the auditory system and brain, and the EEG technique has inherent problems, such as a low SNR of the signal of interest. Therefore in the future non-linear techniques such as artificial neural networks may yield improved performance (e.g., Yang et al (2015)).

Even with perfect reconstruction of the envelope from the EEG, differences between the CT and SRT can still be expected. First of all, the SRT obtained in a behavioral experiment is not infinitely precise, with a typical test-retest difference of around 2 dB. Second, the two measures do not reflect exactly the same thing: the CT presumably reflects relatively early neural coding of the speech envelope, while the SRT is the product of much more extensive processing, including remembering and repeating the sentence. Another difference is procedural in nature: in the behavioral experiment, we collected a response after each sentence was presented, ensuring the subject’s continuous attention. In the EEG experiment we continuously recorded the EEG during the stimulus, and it is likely that the subject’s attention lapsed once in a while. We attempted to mitigate these differences by selecting young, cognitively strong listeners, using low-context speech material, clear instructions, and asking the subjects regular questions during the EEG experiment to ensure they remained attentive.

To translate this method to the clinic, it first needs to be further validated with a more diverse population with a wider age range, including children, various degrees of hearing impairment, different languages, etc., as it is possible that the optimal signal processing parameters depend on these factors (Presacco et al, 2016). It also needs to be investigated to what extent attention influences the results.

### 4.4 Conclusions

There is a missing link between the current behavioral and electrophysiological methods to assess hearing. The behavioral methods can yield a precise measure of speech intelligibility, but suffer from several confounding factors when the goal is to assess how the auditory periphery processes supra-threshold sounds. Current objective methods do not have this confound and can address specific areas in the auditory pathway. However they do not give much insight in how well the patient understands speech due to the use of simple repetitive stimuli. The proposed measure (CT) is based on running speech stimuli and is fully objective. It can on one hand provide valuable information additional to behaviorally measured speech intelligibility in a population where cognitive factors play a role, such as in aging individuals, or during auditory rehabilitation after fitting an auditory prosthesis. On the other hand it enables completely automatic measurement, which is invaluable for testing individuals who cannot provide feedback, for automatic fitting of auditory prostheses, and for closed-loop auditory prostheses that continuously adapt their function to the individual listener in a specific and changing listening environment.

## Acknowledgements

The authors thank Lise Goris and Eline Verschueren for their help with the data acquisition.

## Footnotes

1 http://www.radioboeken.eu/radioboek.php?id=193&lang=NL

## References

Aiken SJ, Picton TW (2008) Human cortical responses to the speech envelope. Ear and Hearing 29(2):139–157

Anderson S, Parbery-Clark A, White-Schwoch T, Kraus N (2013) Auditory brainstem response to complex sounds predicts self-reported speech-in-noise performance. Journal of Speech, Language, and Hearing Research 56(1):31–43

Biesmans W, Das N, Francart T, Bertrand A (2016) Auditory-inspired speech envelope extraction methods for improved eeg-based auditory attention detection in a cocktail party scenario. IEEE Transactions on Neural Systems and Rehabilitation Engineering

Di Liberto GM, O’Sullivan JA, Lalor EC (2015) Low-frequency cortical entrainment to speech reflects phoneme-level processing. Current Biology 25(19):2457–2465

Dillon H (2012) Hearing aids. Thieme, Stuttgart

Ding N, Simon JZ (2011) Neural coding of continuous speech in auditory cortex during monaural and dichotic listening. Journal of Neurophysiology 107(1):78–89

Ding N, Simon JZ (2012) Emergence of neural encoding of auditory objects while listening to competing speakers. Proceedings of the National Academy of Sciences 109(29):1,854–11,859

Ding N, Simon JZ (2013) Adaptive temporal encoding leads to a background-insensitive cortical representation of speech. The Journal of Neuroscience 33(13):5728–5735

Ding N, Simon JZ (2014) Cortical entrainment to continuous speech: functional roles and interpretations. Frontiers in Human Neuroscience 8:311

Ding N, Chatterjee M, Simon JZ (2014) Robust cortical entrainment to the speech envelope relies on the spectro-temporal fine structure. Neuroimage 88:41–46

Doelling KB, Arnal LH, Ghitza O, Poeppel D (2014) Acoustic landmarks drive delta-theta oscillations to enable speech comprehension by facilitating perceptual parsing. Neuroimage 85:761–768

Drullman R, Festen JM, Plomp R (1994a) Effect of reducing slow temporal modulations on speech reception. The Journal of the Acoustical Society of America 95(5):2670–2680

Drullman R, Festen JM, Plomp R (1994b) Effect of temporal envelope smearing on speech reception. The Journal of the Acoustical Society of America 95(2):1053–1064

Edwards E, Chang EF (2013) Syllabic (2-5 hz) and fluctuation (1-10 hz) ranges in speech and auditory processing. Hearing research 305:113–134

Francart T, van Wieringen A, Wouters J (2008) APEX 3: a multi-purpose test platform for auditory psychophysical experiments. Journal of Neuroscience Methods 172(2):283–293

Horton C, Srinivasan R, D’Zmura M (2014) Envelope responses in single-trial eeg indicate attended speaker in a ‘cocktail party’. Journal of neural engineering 11(4):046,015

Hullett PW, Hamilton LS, Mesgarani N, Schreiner CE, Chang EF (2016) Human superior temporal gyrus organization of spectrotemporal modulation tuning derived from speech stimuli. The Journal of Neuroscience 36(6):2014–2026

Kong YY, Somarowthu A, Ding N (2015) Effects. of spectral degradation on attentional modulation of cortical auditory responses to continuous speech. Journal of the Association for Research in Otolaryngology 16(6):783–796

Lalor EC, Pearlmutter BA, Reilly RB, McDarby G, Foxe JJ (2006) The vespa: a method for the rapid estimation of a visual evoked potential. Neuroimage 32(4):1549–1561

Lalor EC, Power AJ, Reilly RB, Foxe JJ (2009) Resolving precise temporal processing properties of the auditory system using continuous stimuli. Journal of neurophysiology 102(1):349–359

Luts H, Jansen S, Dreschler W, Wouters J (2015) Development and normative data for the emish/dutch matrix test. Tech. rep.

McGee TJ, Clemis JD (1980) The approximation of audiometric thresholds by auditory brain stem responses. Otolaryngology-Head and Neck Surgery 88(3):295–303

O’Sullivan JA, Power AJ, Mesgarani N, Rajaram S, Foxe JJ, Shinn-Cunningham BG, Slaney M, Shamma SA, Lalor EC (2014) Attentional selection in a cocktail party environment can be decoded from single-trial eeg. Cerebral Cortex pp 1697–1706

Pasley BN, David SV, Mesgarani N, Flinker A, Shamma SA, Crone NE, Knight RT, Chang EF (2012) Reconstructing speech from human auditory cortex. PLoS Biol 10(1):e1001,251

Peelle JE, Davis MH (2012) Neural Oscillations Carry Speech Rhythm through to Comprehension. Front Psychol 3:320

Picton TW, Dimitrijevic A, Perez-Abalo MC, Van Roon P (2005) Estimating audiometric thresholds using auditory steady-state responses. Journal of the American Academy of Audiology 16(3):140–156

Presacco A, Simon JZ, Anderson S (2016) Evidence of degraded representation of speech in noise, in the aging midbrain and cortex. Journal of neurophysiology 116(5):2346–2355

Shannon RV, Zeng FG, Kamath V, Wygonski J, Ekelid M (1995) Speech recognition with primarily temporal cues. Science 270(5234):303–304

Søndergaard PL, Torrésani B, Balazs P (2012) The Linear Time Frequency Analysis Toolbox. International Journal of Wavelets, Multiresolution Analysis and Information Processing 10(4)

Søndergaard P, Majdak P (2013) The auditory modeling toolbox. In: Blauert J (ed) The Technology of Binaural Listening, Springer, Berlin, Heidelberg, pp 33–56

Woodfield A, Akeroyd MA (2010) The role of segmentation di culties in speech-in-speech understanding in older and hearing-impaired adults. The Journal of the Acoustical Society of America 128(1):EL26–EL31

Yang M, Sheth SA, Schevon CA, Gmm II, Mesgarani N (2015) Speech reconstruction from human auditory cortex with deep neural networks. In: Sixteenth Annual Conference of the International Speech Communication Association

